# *Pseudomonas syringae* Increases Water Availability in Leaf Microenvironments via Production of Hygroscopic Syringafactin

**DOI:** 10.1101/626630

**Authors:** Monica N. Hernandez, Steven E. Lindow

## Abstract

The epiphytic bacterium *Pseudomonas syringae* strain B728a produces the biosurfactant syringafactin which is hygroscopic. The water absorbing potential of syringafactin is high. At high relative humidities, syringafactin attracts 250% of its weight in water but is less hygroscopic at lower relative humidities. This suggests that syringafactin’s benefit to the producing cells is strongly context-dependent. The contribution of syringafactin to the water availability around cells on different matrices was assessed by examining water availability biosensor strains that express *gfp* via the water-stress activated *proU* promoter. Wild-type cells exhibited significantly less GFP fluorescence than a syringafactin-deficient strain, on humid but dry filters as well as on leaf surfaces indicating higher water availability. When infiltrated into the leaf apoplast, wild-type cells also subsequently exhibited less GFP fluorescence than a syringafactin-deficient strain. These results suggest that the apoplast is a dry, but humid environment and that, just as on dry but humid leaf surfaces, syringafactin increases liquid water availability and reduces the water stress experienced by *P. syringae*.

**IMPORTANCE:** Many microorganisms, including the plant pathogen *Pseudomonas syringae*, produce amphiphilic compounds known as biosurfactants. While biosurfactants are known to disperse hydrophobic compounds and reduce water tension, they have other properties that can benefit the cells that produce them. Leaf colonizing bacteria experience frequent water stress since liquid water is only transiently present on or in leaf sites that they colonize. The demonstration that syringafactin, a biosurfactant produced by *P. syringae*, is sufficiently hygroscopic to increase water availability to cells, thus relieving water stress, reveals that *P. syringae* can modify its local habitat both on leaf surfaces and in the leaf apoplast. Such habitat modification may be a common role for biosurfactants produced by other bacterial species that colonize habitats that are not always water saturated such as soil.

## INTRODUCTION

While leaf surfaces support large numbers of bacteria, leaves are considered a relatively harsh environment for bacterial colonization. Leaves are frequently dry environmental habitats that are also subject to high ultraviolet fluxes as well as fluctuations in temperature and humidity (1, 2, 3, 4, 5). Desiccation stress is considered one of the major factors limiting bacterial survival on leaves (6, 7). As in many environments, the frequent lack of water on leaves is expected to limit the ability of epiphytes like *Pseudomonas syringae* to colonize leaf surfaces. However, *P. syringae* successfully colonizes and survives on the surfaces of leaves, often subsequently causing disease in its host plant after it enters the leaf interior (4). The leaf apoplast is comprised largely of air-filled voids between parenchymal cells that facilitate gas exchange for photosynthesis making it a humid, but dry environment (4, 8). Transcriptomic analysis of *P. syringae* recovered from both epiphytic and endophytic sites reveals the high expression of genes involved in tolerance of water stress (9). This supports the model that water limitation is experienced by this species both in the interior and exterior of leaves. However, the traits that enable *P. syringae* to grow and survive on and in dry leaves are poorly understood.

While leaves are frequently dry, the relative humidity (RH) of air surrounding the leaf surface is expected to often differ substantially from that of the air surrounding the leaf. Because of friction with the leaf surface, air movement is rapidly inhibited as it crosses the leaf creating a thin layer of still air known as the laminar boundary layer that surrounds the leaf. The thickness of this layer is inversely proportional to wind speed but is usually less than about 10 mm (10). Much of the water vapor that exits the leaf via its stomata is apparently retained within the laminar boundary layer (2, 3, 6, 11, 12, 13, 14, 15, 16). Thus, the air surrounding the leaf surface can exhibit a much higher RH than that in the atmosphere away from the leaf (10, 17). While the high RH expected in a boundary layer would reduce the rate of evaporation of water from bacterial cells on the leaf surface, it would not be expected to eliminate the osmotic or matric water stresses that cells would experience when liquid water is not present.

Biosurfactants, are amphiphilic compounds produced by various microorganisms (18). Most studies of biosurfactants have described their ability to disperse hydrophobic compounds, often enabling their consumption by the biosurfactant-producing bacteria (18, 19). Most biosurfactants can also reduce water tension, thereby enabling the dispersal of water across hydrophobic surfaces such as leaves (3, 20). This trait might be beneficial to epiphytic bacteria. Burch *et al*. (2) recently reported that certain biosurfactants such as syringafactin, a biosurfactant produced by *P. syringae*, also have the under-appreciated characteristic of being hygroscopic. Syringafactin is a lipopeptide containing eight amino acids linked to an acyl tail, making it amphipathic (21). The peptide head of this molecule contains several hydroxyl groups capable of hydrogen bonding with water. This structure suggested that syringafactin could interact with and absorb water. Burch *et al.* (2) verified the hygroscopic nature of syringafactin by showing that, after being desiccated, syringafactin could be rewetted when exposed to a water-saturated atmosphere. Syringafactin production appears to be beneficial to *P. syringae* on leaf surfaces. When a wild-type *P. syringae* strain and a *syfA* mutant strain deficient in syringafactin production were co-inoculated onto bean leaves in a field experiment, the wild-type strain maintained greater population sizes on plants than the *syfA* mutant strain (2). This suggested that the wild-type strain was more tolerant of desiccation stresses experienced during fluctuating environmental conditions in the field.

The goal of this study was to test the hypothesis that the contributions of syringafactin to the epiphytic fitness of *P. syringae* is due to its ability to absorb water vapor from the air, thereby maintaining a more hydrated state in the vicinity of cells producing this compound. This would reduce the cells’ water stress while on plants in the absence of liquid water. Such a role would require syringafactin to bind abundant water under the conditions that cells would experience on leaf surfaces. While the RH experienced by bacteria on the surface of plants is expected to be higher than that in air surrounding the plant, the actual RH and its temporal variability on leaves is unknown. Furthermore, although syringafactin was shown to bind abundant water in a water-saturated environment (2) it is not clear whether it can do so under conditions experienced by cells on a plant. By determining both the RH-dependent water-binding capabilities of syringafactin as well as the apparent water status of bacteria on leaves, it should be possible to both test the hypothesis above and provide insight as to the water environment experienced by cells on leaves. Such information is needed to determine under what contexts syringafactin would benefit the producing cells. To measure the water status of cells, we utilize a whole-cell bacterial biosensor described by Axtell and Beattie (1) that assesses the expression of *proU*, a gene contributing to production of the compatible solute proline, by linking it to a GFP reporter gene. Cells harboring this reporter gene construct exhibit GFP fluorescence that is directly proportional to the level of either the matric or osmotic stress that they experience (1). In this study, we assessed the water availability experienced by both a wild-type *P. syringae* B728a strain and a *syfA* mutant strain deficient in syringafactin production on both the surface and interior of plants by quantifying the fluorescence of individual bacterial cells using epifluorescence microscopy. As described below, our results strongly suggest that syringafactin production by *P. syringae* can reduce the desiccation stress that cells experience both on the leaf surface and in the leaf apoplast. This work thus reveals an important and previously unrecognized role for microbial biosurfactants in the varied environments that such epiphytes colonize.

## RESULTS

### Syringafactin is very hygroscopic only at high relative humidities

Since the hypothesized ecological role of syringafactin depends on its ability to interact with water, we examined the conditions over which syringafactin would absorb water. Purified dehydrated syringafactin was exposed to different controlled RH conditions maintained by suspension over different saturated salt solutions (Fig. 1). The weight of the syringafactin was determined both before exposure and after 3 days of exposure to a given RH. Although the water absorbing potential of syringafactin generally increased with increasing RH, it absorbed less than its own weight in water over most levels of atmospheric water saturation. Importantly, its water-binding capacity increased dramatically at relative humidities greater than about 97%; however, it absorbed 250% its weight in water in fully water-saturated air (Fig. 1). This indicates that syringafactin is especially hygroscopic at high RH and that its maximum potential ecological value may be in conditions of high levels of atmospheric water saturation.

**FIG 1.**
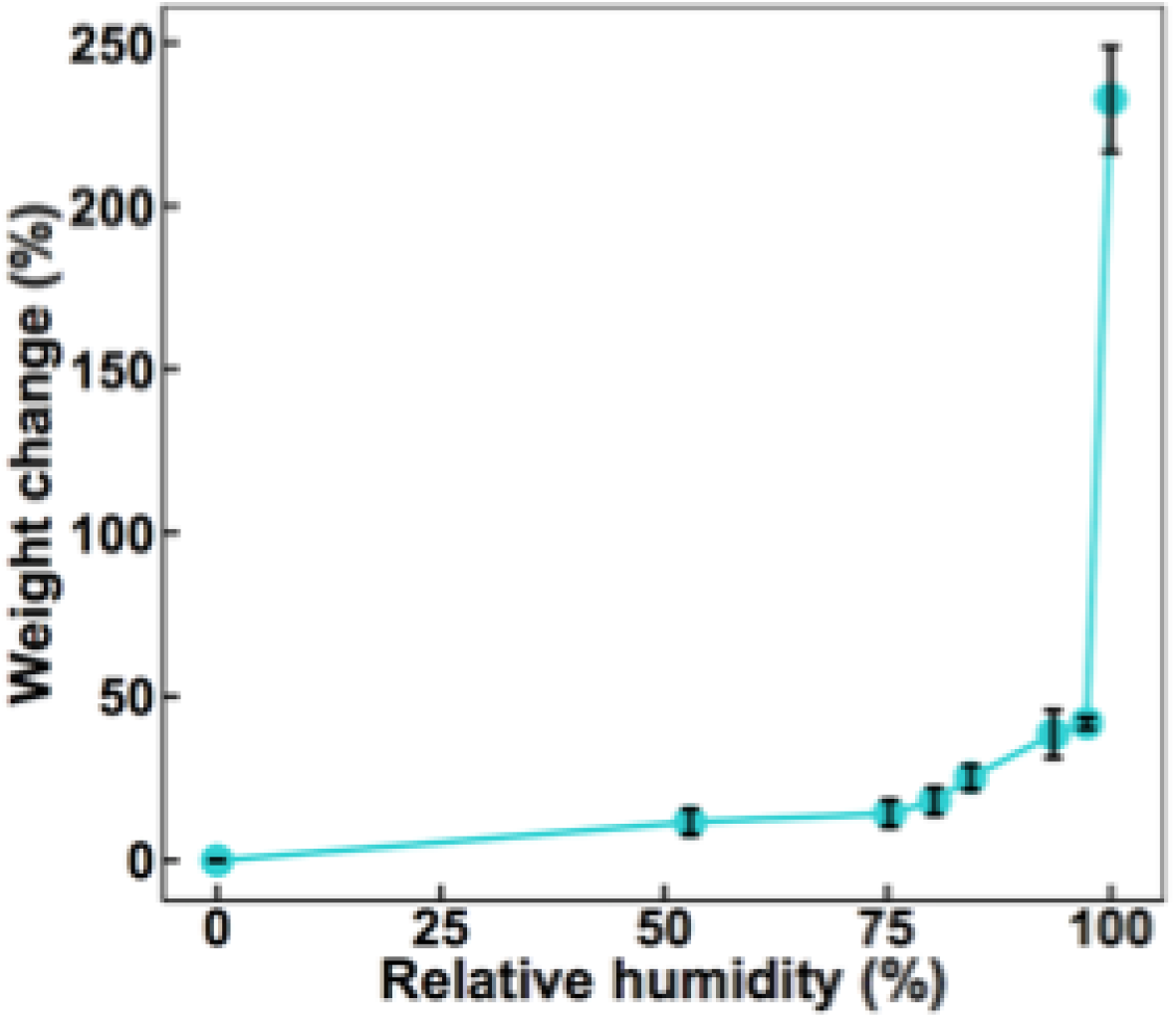
Water binding by syringafactin. Weight gain due to water absorption, expressed as a proportion of the initial weight of dehydrated syringafactin exposed for 3 days to atmospheres containing the relative humidity shown on the abscissa. The vertical bars represent the standard error of the mean percentage weight change.

### Syringafactin contributes to water availability to cells on filters

To determine whether syringafactin made enough water available to bacterial cells to alleviate water stress, we compared the water stress exhibited by *P. syringae* cells differing in syringafactin production when immobilized on membrane filters. Since they would lack the humid laminar boundary layer of leaves, filters were used as a more direct means to determine the conditions under which, and the degree to which, the water status of cells was modulated by the presence of syringafactin. Cells of the wild-type and *syfA* mutant strain harboring the *proU:gfp* reporter gene construct grown in a low salt minimal medium, in which they would be expected to experience the same low level of water stress, did not differ in their expression of GFP fluorescence before application to filters (Fig. 2). Syringafactin production therefore does not influence the cellular response to water availability. The wild-type and a *syfA* mutant *P. syringae* strain were then grown on filters placed on agar plates for 8 h before the filters were transferred to chambers maintaining 52% RH or 100% RH. Filters were incubated in the chambers for 4 h and then immersed in a low salt-containing minimal nutrient medium for 2 h to resuscitate cells and enable the translation of GFP resulting from the transcription of the reporter gene. As a positive control, exogenous syringafactin was added to cells of the *syfA* mutant strain after they had grown on the filter. Wild-type cells exhibited significantly less GFP fluorescence than the *syfA* mutant when incubated at 52% RH (Fig. 3A). The GFP fluorescence of the *syfA* mutant strain exposed to exogenous syringafactin was similar to the wild-type strain (Fig. 3A). Similarly, at 100% RH, the GFP fluorescence exhibited by the *syfA* mutant strain to which exogenous syringafactin had been applied was significantly less than that exhibited by this strain in the absence of added syringafactin (Fig. 3B). Furthermore, the GFP fluorescence of the *syfA* mutant was still higher than that of the wild-type strain when both were incubated at 100% RH (Fig. 3B). The finding that both the wild-type strain alone and the *syfA* mutant strain with applied syringafactin exhibited similarly lower GFP fluorescence than the *syfA* mutant strain itself at 100% RH supports our hypothesis that syringafactin is not only capable of making water more available to cells under high RH conditions, but that wild-type cells produce sufficient amounts of syringafactin to confer this phenotype. Surprisingly, this trend was also seen at 52% RH, suggesting that the lesser amounts of water retained by syringafactin at this lower RH still was enough to reduce somewhat the water stress that producing cells experienced.

**FIG 2.**
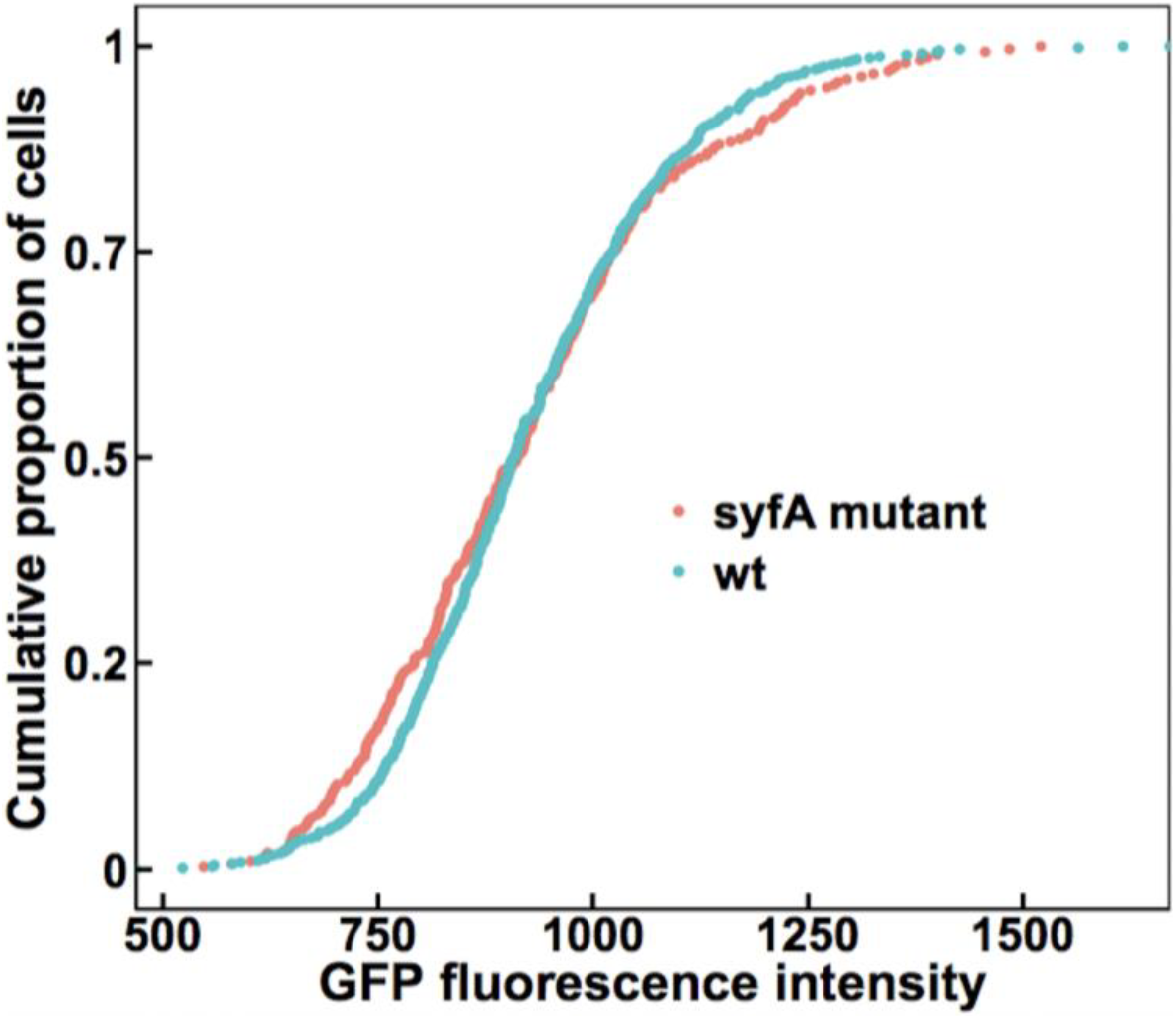
In liquid media, syringafactin production does not affect water stress of *P. syringae*. Single-cell GFP fluorescence exhibited by wild-type *P. syringae* strain B728a (blue) or a *syfA* mutant strain (red) harboring a *proU:gfp* reporter when grown on half-strength 1/2-21C minimal media plates. The mean fluorescence of the wild-type cells was 926 (n = 577) while that of the *syfA* mutant cells was 925 (n = 301). Single-cell fluorescence was quantified by microscopy. The two distributions do not differ significantly by the two-sample t-test (p-value = 0.68).

**FIG 3.**
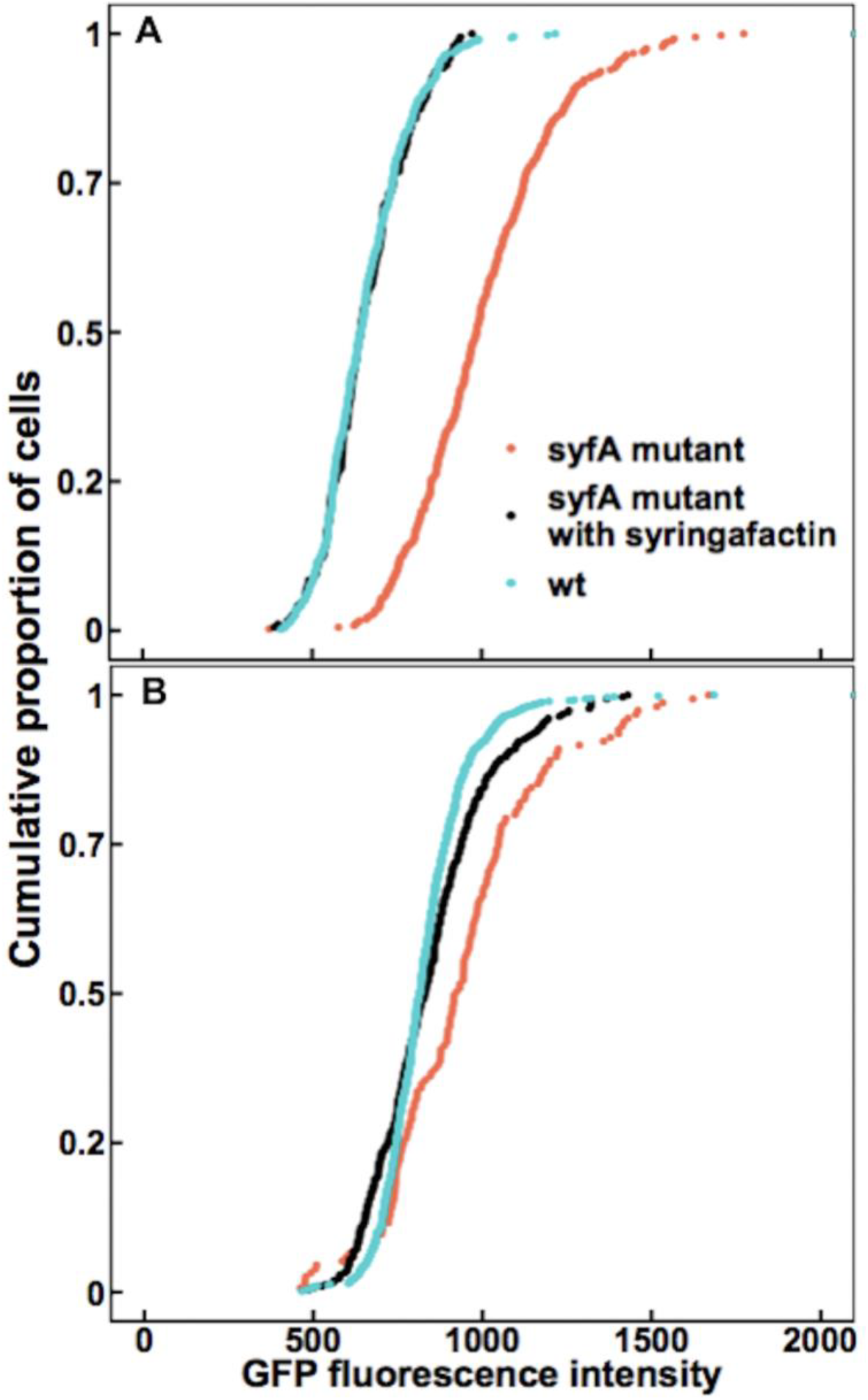
Syringafactin reduces water stress experienced by *P. syringae* on filters. Lower water stress experienced by wild-type *P. syringae* cells on filters at (A) 52% RH and (B) 100% RH than *syfA* mutant cells. Wild-type (blue) and a *syfA* mutant strain (red) harboring a *proU:gfp* reporter as well as the *syfA* mutant with added syringafactin (black) were incubated on membrane filters on minimal medium plates for 8 h before filters were transferred to glass slides placed in chambers that maintained either 52% RH (A) or 100% RH (B). Single-cell fluorescence was quantified by microscopy. (A) After exposure to 52% RH the median fluorescence of wild-type cells was 643 (n = 449) while that of *syfA* mutant cells was 984 (n = 366) and that of *syfA* mutant cells with added syringafactin was 643 (n = 192). Significance between sample distributions was tested using the Wilcoxon rank-sum test (wt vs *syfA* mutant: p-value < 2.2 × 10^−16^, W = 153,460; wt vs *syfA* mutant with syringafactin: p = 0.70, W = 43,693; *syfA* mutant vs *syfA* mutant with syringafactin: p-value < 2.2 × 10^−16^, W = 65,276). (B) After exposure to 100% RH the median fluorescence of wild-type cells was 815 (n = 767) while that of *syfA* mutant cells was 924 (n = 155) and that of *syfA* mutant cells with syringafactin was 831 (n = 308). Significance between sample distributions was tested using the Wilcoxon rank-sum test (wt vs *syfA* mutant: p-value = 7.71 × 10^−11^, W = 39,768; wt vs *syfA* mutant with syringafactin: p-value = 0.36, W = 113,890; *syfA* mutant vs *syfA* mutant with syringafactin: p-value = 2.78 × 10^−6^, W = 30,238).

### Syringafactin improves water availability to cells on leaf surfaces irrespective of the dryness of air away from the leaf

Given that syringafactin could make water more available to cells on abiotic surfaces, we determined the extent of water stress experienced by bacterial cells on leaves exposed to various environmental conditions and asked whether cells could ameliorate this stress by producing syringafactin. We hypothesized that at a high RH the wild-type cells would exhibit lower GFP fluorescence than the *syfA* mutant cells due to the attraction of water by syringfactin production. To test this, wild-type and *syfA* mutant strains harboring the *proU:gfp* reporter gene fusion were sprayed onto the leaves of bean plants that were then immediately placed in a 100% RH chamber for 2 days to enable bacterial growth and production of any extracellular products. The sprayed leaves initially were covered with many small droplets of bacterial suspension, but after 2 days, most of the leaf was free of any droplets. Instead, only a few water droplets persisted on the leaves, suggesting that the water had evaporated from leaves to condense on the chamber walls and any residual water was also redistributed to produce large dry areas on the leaf surface. When examined 2 days after inoculation, wild-type cells exhibited less GFP expression than the *syfA* mutant cells (Fig. 4). Presumably, many of the cells on these leaves were localized at sites on the leaf that were devoid of liquid water and thus experienced water stress although the humidity on the leaves must have been near 100% RH.

**FIG 4.**
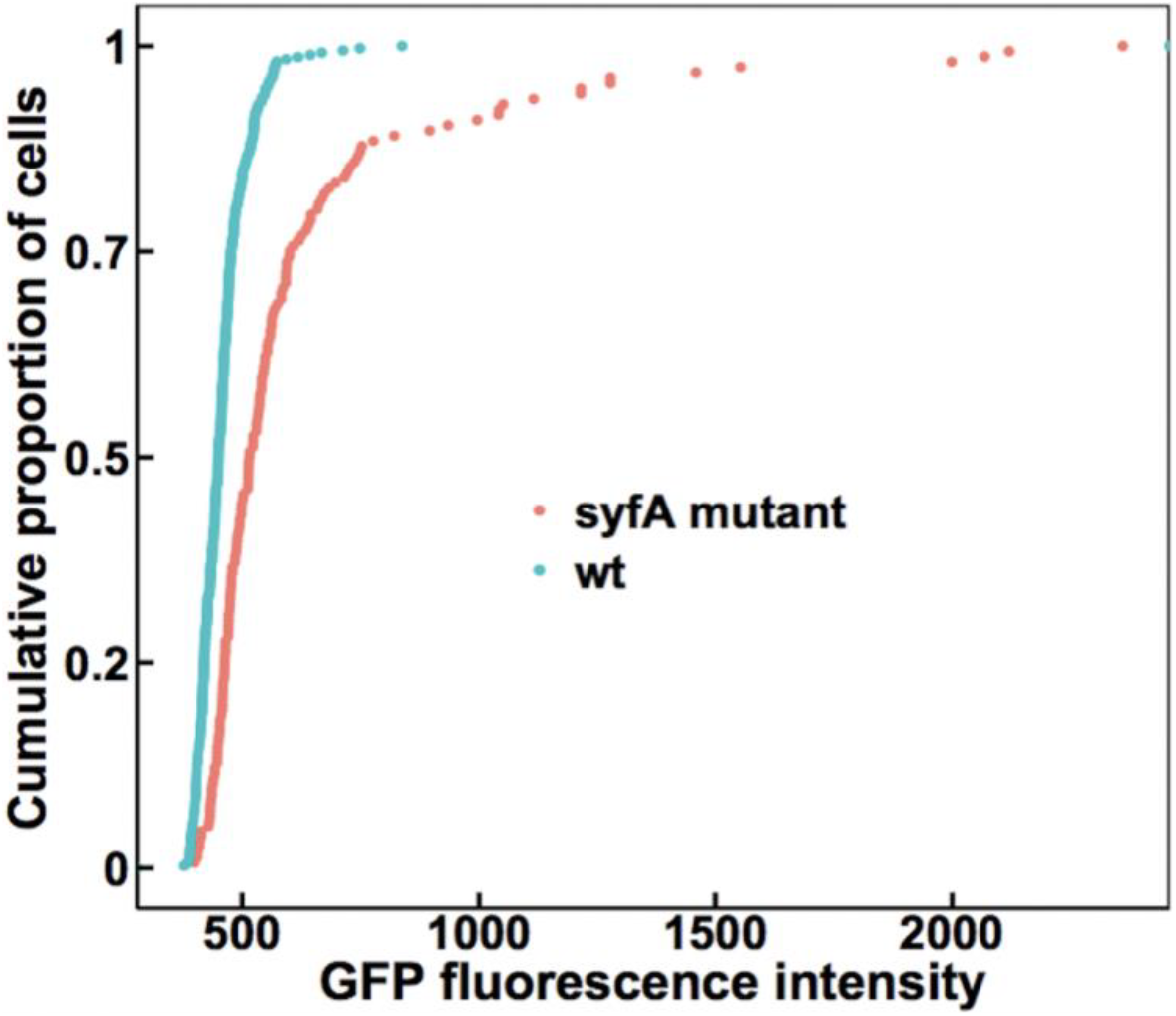
Syringafactin-producing strains of *P. syringae* experience more water availability than *syfA* mutants on humid leaves. GFP fluorescence exhibited by wild-type *P. syringae* strain B728a (blue) or a *syfA* mutant strain (red) harboring a *proU:gfp* reporter when recovered from the leaves of plants incubated at 100 % RH for 2 days. The median fluorescence of wild-type cells was 450 (n = 371) while that of *syfA* mutant cells was 516 (n = 156). Single-cell fluorescence was quantified by microscopy. Significance between sample distributions was tested using the Wilcoxon rank-sum test (wt vs *syfA* mutant: W = 45,442, p-value < 2.2 × 10^−16^).

To determine the environmental contexts under which syringafactin could be produced and under which it could confer protection against desiccation stress, we exposed cells of *P. syringae* to various drying conditions on leaves. Leaves were exposed sequentially to wet conditions in which bacterial cells could multiply on leaves, followed by dry conditions where multiplication ceases. Though a difference in GFP fluorescence was observed between the wild-type and *syfA* mutant strains harboring the *proU:gfp* reporter gene when cells were applied to dry plants that remained dry throughout the experiments (Fig. S1), it is noteworthy that this difference seemed to be caused by only about 20% of the cell population rather than a majority. Indeed, in a subsequent experiment, most of the cells died quickly after immediate drying on surfaces (Fig. S2). This phenomenon has been seen previously on leaves (7). Therefore, these cells would not have been capable of expressing the reporter gene. We therefore assessed the water availability to bacteria exposed to dry conditions that followed moist incubation conditions after migration of cells to the plants which enabled bacterial colonization and any habitat modification to occur. In one such condition, inoculated bean plants were immediately incubated in a 100% RH chamber for 2 days before leaf surfaces were dried by exposure of the plants to 50% RH for 20 min and then incubated at 100% RH for 2 more hours (Fig. 5). Such plants would have experienced liquid water on leaf surfaces only in the initial 2 days incubation period, and cells would have subsequently found themselves on dry leaves that were exposed to varying RH conditions. We presume that the apparent transcription of the *proU:gfp* reporter gene, as evidenced by the GFP fluorescence output, would have reflected the conditions experienced by the cells on the final 2 h dry but humid incubation period. Under these conditions it was evident that while the wild-type strain had exhibited the same relatively low GFP fluorescence indicative of low water stress as it had on leaves continually exposed to high RH conditions, the *syfA* mutant strain apparently experienced considerable water stress as indicated by its high GFP fluorescence (Fig. 5). Given that the GFP fluorescence of most *syfA* mutant cells was much higher than that of the wild-type strain, it suggested that they experienced more water stress than the wild-type strain (Fig. 5). These results suggest that syringafactin production by the wild-type strain could sequester sufficient water to avoid desiccation stress when cells exposed to desiccating conditions were subsequently exposed to a water-saturated environment.

**FIG 5.**
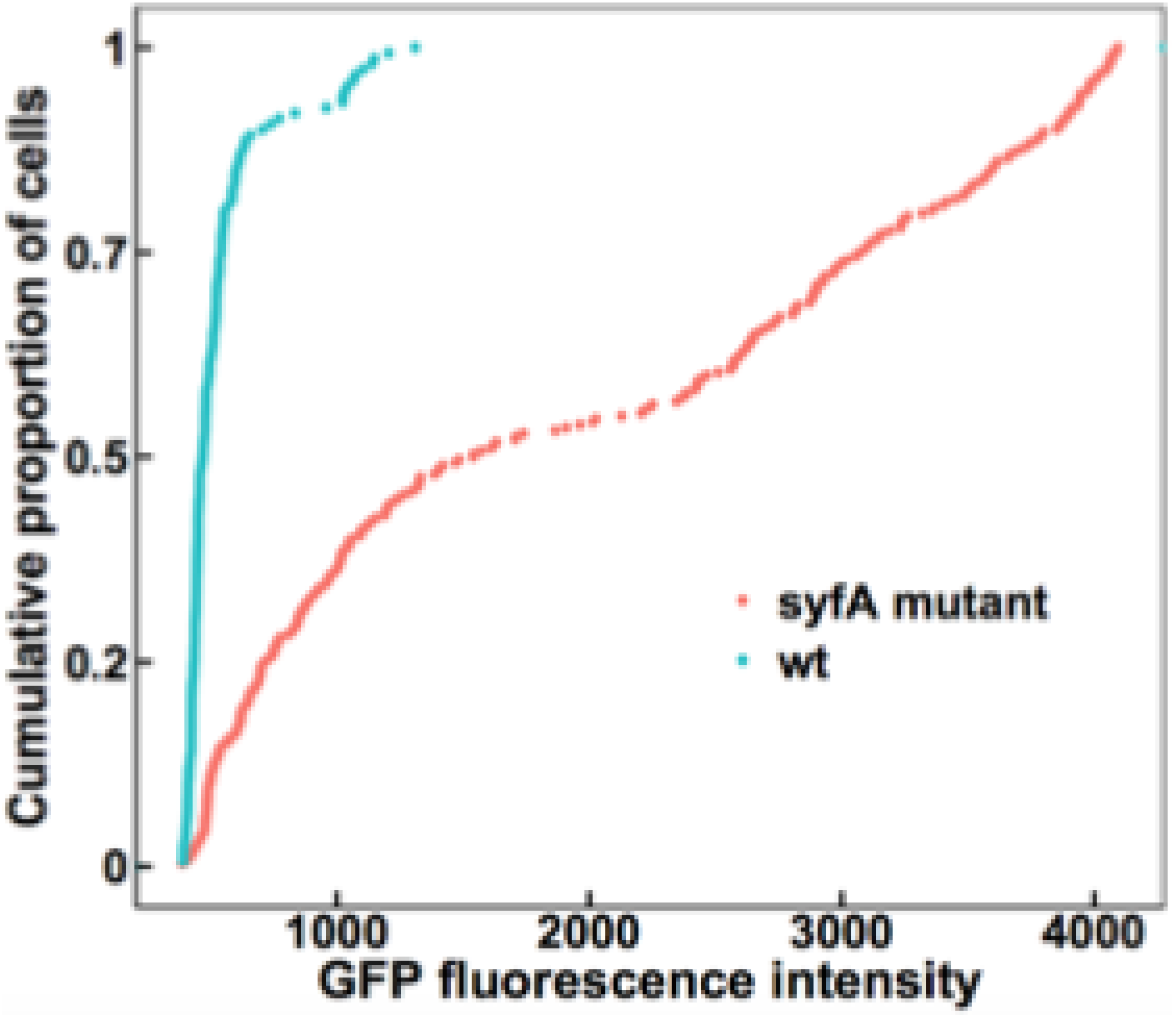
Wild-type cells experience more water availability than *syfA* mutant cells after transient exposure to low RH on plants. GFP fluorescence exhibited by wild-type *P. syringae* strain B728a (blue) or a *syfA* mutant strain (red) harboring a *proU:gfp* reporter when recovered from the leaves of plants incubated at 100 % RH for 2 days followed by drying at 50% RH for 20 min before re-exposure to 100% RH for 2 h. The median fluorescence of the wild-type cells was 467 (n = 150) while that of the *syfA* mutant cells was 1,541 (n = 282). Single-cell fluorescence was quantified by microscopy. Significance between sample distributions was tested using the Wilcoxon rank-sum test (wt vs *syfA* mutant: W = 38,207, p-value < 2.2 × 10^−16^).

### Wild-type cells experience less desiccation stress than *syfA* mutant cells when exposed to fluctuating RH conditions at less than full atmospheric water saturation

Given that plants frequently experience conditions of less than full atmospheric water saturation (100% RH) under field conditions (3), we examined the potential for syringafactin to modulate the water availability of cells on the surface of leaves under these conditions. Bean leaves, sprayed with either the wild-type or the *syfA* mutant strain harboring the *proU:gfp* reporter gene fusion, were immediately incubated at 100% RH for 2 days to enable bacterial growth and metabolism. The plants were then subsequently exposed to 50% RH for 1 h to allow liquid water to evaporate from the leaf before being incubated at 97% RH for 2 more days. As was seen when such dried, colonized leaves were subsequently exposed to 100% RH, the GFP fluorescence of the *syfA* mutant strain was significantly higher than that of the wild-type strain (Fig. 6) indicating that it exhibited a higher degree of water stress than the wild-type strain. Given that the water-binding capability of syringafactin at 97% RH is much lower than that in a fully water-saturated atmosphere, it seems likely that the RH experienced by cells on plants incubated at 97% RH was actually much higher because of the modulation of the air in the laminar boundary layer surrounding leaves by water vapor released by plant transpiration. This would enable syringafactin to bind water and thus hydrate the cells of the syringafactin-producing strain.

**FIG 6.**
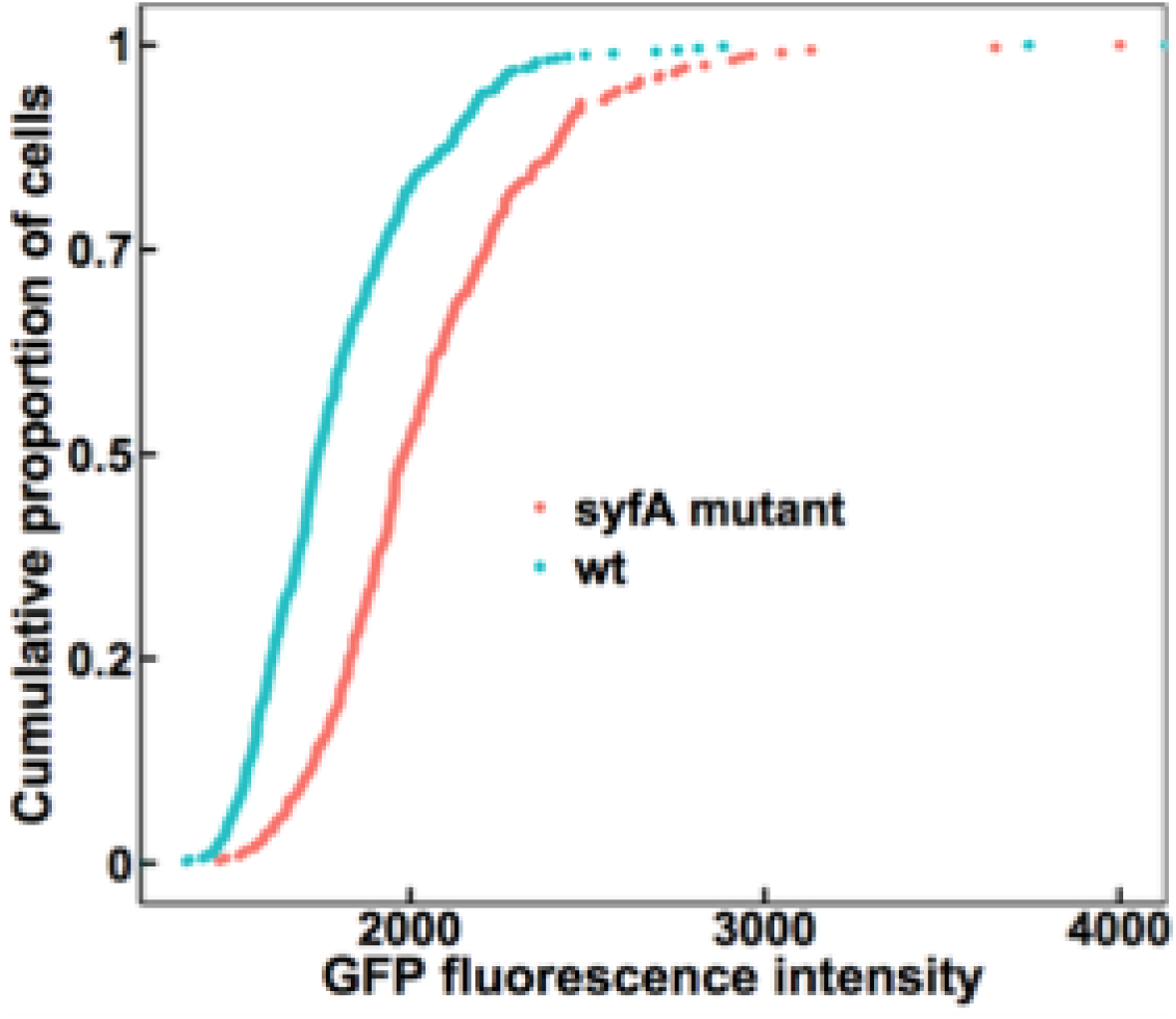
Wild-type cells experience more water availability than *syfA* mutant cells when exposed to 97% RH on leaf surfaces. GFP fluorescence exhibited by wild-type *P. syringae* strain B728a (blue) or a *syfA* mutant strain (red) harboring a *proU:gfp* reporter when recovered from the leaves of plants incubated at 100 % RH for 2 days followed by drying at 50% RH for 20 min before re-exposure to 97% RH for 2 days. The median fluorescence of the wild-type cells was 1,741 (n = 489) while that of the *syfA* mutant cells was 1,984 (n = 324). Single-cell fluorescence was quantified by microscopy. Significance between sample distributions was tested using the Wilcoxon rank-sum test (wt vs *syfA* mutant: W = 119,610, p-value < 2.2 × 10^−16^).

### Syringafactin helps make water available to bacteria in the leaf apoplast

After colonizing the leaf surface, cells of *P. syringae* can enter the leaf apoplast through stomata where they can grow to sufficient numbers to eventually cause disease (4). Given that the apoplast is often dry, at least initially after bacterial entry, we determined if syringafactin production could reduce water stress in this habitat just as it apparently does on the leaf surface. Wild-type and *syfA* mutant cells harboring the *proU:gfp* reporter gene construct were infiltrated under vacuum into bean leaves. Before infiltrating the cells into the leaves, aliquots of cells taken from the liquid culture used as inoculum were assessed for GFP fluorescence to determine if the two strains differed in apparent water availability at time 0. While the GFP fluorescence exhibited by the two strains was initially quite similar (Fig. 7A), when assessed 24 h after inoculation, cells of the *syfA* mutant strain exhibited higher GFP fluorescence than that of cells in the wild-type strain, indicating that they were beginning to experience somewhat more desiccation stress than the wild-type strain (Fig. 7B). By 48 h after inoculation, the GFP fluorescence of both strains in the apoplast was much higher than after 24 h, indicating that the intercellular spaces had become even drier during the infection process (Fig. 7C). Importantly, by 48 h after incubation, *syfA* mutant cells exhibited much higher GFP fluorescence than the wild-type cells indicating that they experienced a higher water stress than that of the wild-type strain (Fig. 7C). Thus, syringafactin production by the wild-type strain had ameliorated the water stress that it otherwise would have experienced.

**FIG 7.**
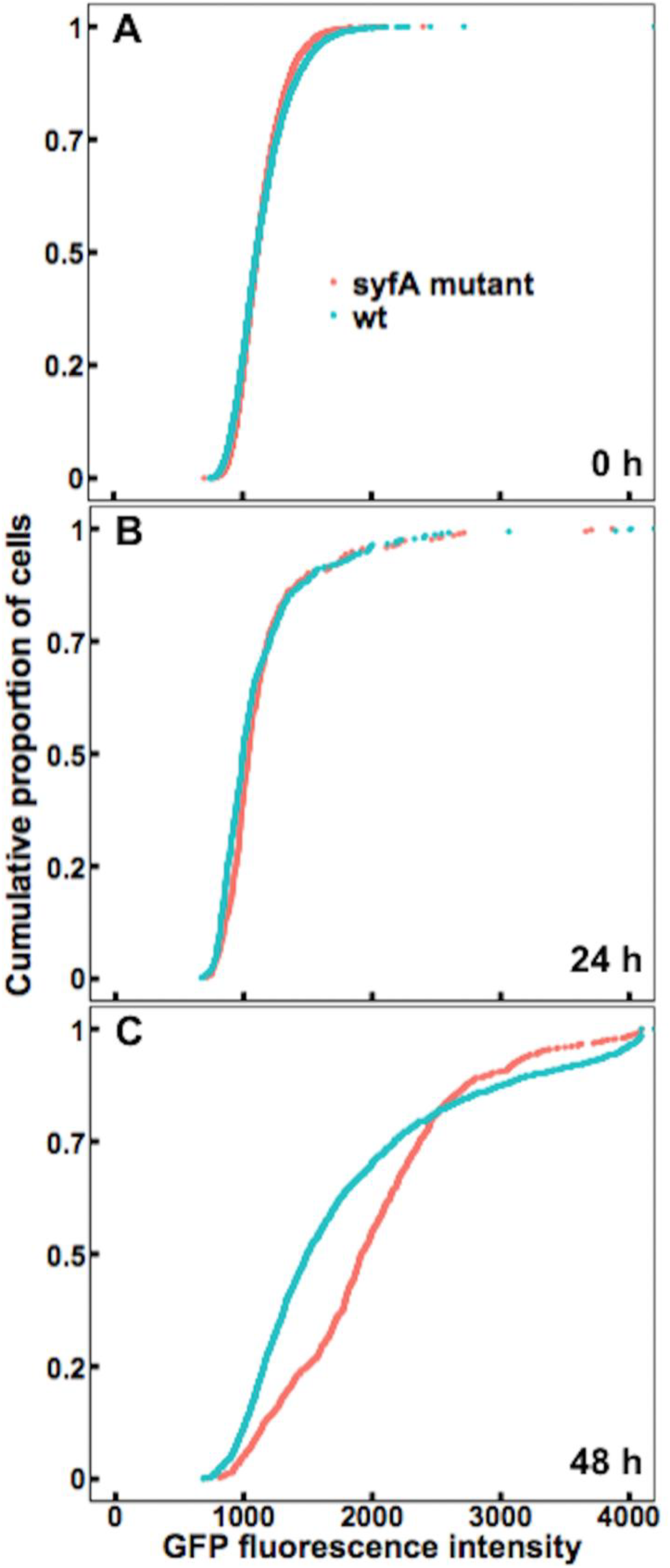
Wild-type *P. syringae* cells in the leaf apoplast experience more water availability than a *syfA* mutant. GFP fluorescence exhibited by wild-type strain B728a (blue) or a *syfA* mutant strain (red) harboring a *proU:gfp* reporter when recovered from the interior of plants 0 h (A) 24 h (B) or 48 h (C) after cells were vacuum infiltrated into the apoplasts of bean leaves. (A) After 0 h, the median fluorescence of the wild-type cells was 1,107 (n = 4,023) while that of the *syfA* mutant cells was 1,112 (n = 4,000). (B) After 24 h, the median fluorescence of the wild-type cells was 992 (n = 447) while that of the *syfA* mutant cells was 1,035 (n = 336). (C) After 48h, the median fluorescence of the wild-type cells was 1,492 (n = 1,528) while that of the *syfA* mutant cells was 1,914 (n = 908). Single-cell fluorescence was quantified by microscopy. Significance between sample distributions was tested using (A) the two-sample t-test (wt vs *syfA* mutant: degrees of freedom = 8,021, p-value = 0.97); (B) Wilcoxon rank-sum test (wt vs *syfA* mutant: W = 83,004, p-value = 0.01); (C) Wilcoxon rank-sum test (wt vs *syfA* mutant: W = 856,960, p-value < 2.2 × 10^−16^).

## DISCUSSION

This study determines whether syringafactin plays an important role in the epiphytic and endophytic growth of *P. syringae*. It is reasonable to assume that liquid water is required for bacterial growth in and on leaves, since it would be required to mobilize soluble nutrients (8). However, leaf surfaces are dry much of the time and cells must survive such conditions in order to grow during the brief periods when water might become available. Thus, life on a leaf surface is probably stressful, since it is a dry environment that is only transiently wet (1, 2, 5, 6, 22).

We found that the hygroscopic biosurfactant syringafactin produced by *P. syringae* B728a reduces the water stress experienced by this species both on dry leaves and in the apoplast by attracting water vapor from the atmosphere and also perhaps by retaining water after cells have been wetted. Syringafactin is quite hygroscopic under the high RH conditions expected in both the apoplast and the humid laminar boundary layer immediately above leaf surfaces. Although syringafactin can bind substantial water only in air that is nearly fully saturated with water vapor (Fig. 1), these conditions are consistent with models of the abiotic conditions that prevail on leaves. Such models predict that the water content of air immediately surrounding the leaf, known as the laminar boundary layer, can differ greatly from that of air further away from the leaf (10, 11, 15, 23) since it traps water vapor exiting the leaf via stomata. Thus, a very humid microenvironment is proposed to surround even dry leaves (2, 3, 6, 11, 12, 13, 14, 15, 16). While syringafactin absorbs substantial amounts of water only at a RH greater than 97%, cells that exist within a laminar boundary layer apparently often experience such a high RH. Even when plants are exposed to a relatively dry environment (97% RH), the reduced water stress exhibited by the wild-type strain compared to that of the *syfA* mutant strain on such dry leaves (Fig. 6) can be attributed to water made available by binding to the syringafactin produced by the wild-type strain. Thus, even though the water-binding capability of syringafactin is strongly dependent on the RH of the atmosphere, cells in and on plants apparently reside in a sufficiently humid atmosphere for them to benefit from water binding by syringafactin.

This study also supports predictions that the physical characteristics of leaves lead to great spatial heterogeneity in the microhabitats that cells experience. For example, the *syfA* mutant strain harboring the *proU:gfp* reporter gene exhibited greater GFP fluorescence than the wild-type strain even on plants that had been maintained in a water-saturated environment after they were sprayed with bacterial suspensions (Fig. 4). These results suggest that many cells of this mutant strain experienced lesser amounts of water than those of the wild-type strain. The observation that water limitations might occur under such a scenario supports and advances previous models of the movement and distribution of water on leaves. For instance, rather than being evenly distributed across the leaf surface, water is thought to be more prevalent at the base of trichomes or at leaf veins or cracks in the cuticle (6, 7, 24). Nutrients are also apparently very spatially variable in their abundance (25) but their coincidence with sites where water is most likely to be retained is unknown. While there was apparently a very slow removal of water from the leaf surface as a whole even at 100% RH conditions in our study, the remaining water also seemed to have been redistributed such as in other studies, collecting in a few sites on the leaf (6, 7, 24). This is quite apparent in this study since leaves that were sprayed with cells initially harbored many small droplets, but after 2 days in a water-saturated environment leaves contained only a few apparent water droplets. Most of the leaf surface was apparently devoid of water and none of the leaves harbored films of water. These observations along with our results further support the hypothesis that the hydrophobic leaf surface is very poorly wettable, and that cells migrating to most areas of the leaf would experience a relatively dry environment due to the localized retention of water even under relatively humid conditions - and would often encounter a complete lack of water (1, 2, 3).

The hypothesis that syringafactin production benefited cells by making water more available, even in leaves exposed to relatively low RH, was supported by the results observed both in cells on continuously wet and humid leaves (Fig. 4) and when leaf surfaces were dried before again being placed in humid conditions (Fig. 5). In both cases, the wild-type strain presumably could have made syringafactin under the moist conditions initially present on leaves after inoculation. As expected, a much higher level of GFP expression was seen in the *syfA* mutant strain when plants were exposed to 50% RH rather than when plants were in a continually water-saturated environment. Given that a quantitative relationship between GFP expression of cells harboring such a reporter gene construct and the level of either matric or osmotic stress to which cells were exposed to has been observed (1), it seems clear that many of the cells of the *syfA* mutant experienced lower water availability on these drier leaves than those on the plants maintained under humid conditions. Importantly, a much larger difference in GFP fluorescence exhibited by the wild-type and *syfA* mutant strains was observed when cells were inoculated on plants exposed to low humidity conditions after their growth on the plant (Fig. 5). We presume that any liquid water would have been removed from both wild-type and *syfA* mutant strains in such a strong drying event, but apparently only the wild-type strain could become rehydrated or would have locally retained water due to its production of the hygroscopic syringafactin. Additional support for this model was also provided by studies in which plants were incubated at 97% RH after colonization. Under these conditions the wild-type strain still exhibited less GFP fluorescence compared to the *syfA* mutant strain indicating that it was wetter (Fig. 6). These results suggest that after being produced during periods of metabolic activity of cells on leaves, syringafactin benefits cells during their subsequent, and probably inevitable, exposure to periods of low RH by enabling the rehydration or suppressing the dehydration of cells. As discussed above, the RH of the air at the leaf surface is probably often above 97% in the plants incubated in a chamber at such a humidity. In this setting, the wild-type cells would be expected to be more highly hydrated than mutants lacking the ability to have produced syringafactin. Such a scenario is supported by the results of Burch *et al*. (2) who observed that even though the ability of syringafactin to wet an abiotic surface was lost at 50% RH, when the filter was re-exposed to 100% RH the syringafactin was able to rehydrate and rewet the surface. This suggests that once cells have grown on moist leaves and produced syringafactin, they subsequently benefit from its production by being able to absorb water and make it more available to cells during fluctuating atmospheric moisture conditions.

Features of syringafactin suggest that it might only influence the environment of cells very locally. While it is highly hygroscopic, syringafactin is apparently not readily dispersed across the leaf surface since over 70% of purified syringafactin was bound to the waxy cuticle of the leaf after topical application (2). This observation suggested that syringafactin largely remains in the local environment of the bacterium that secreted it. This hypothesis was further supported by the observation that the *syfA* mutant strain inoculated onto bean plants did not maintain epiphytic population sizes as large as the wild-type strain irrespective of whether it was inoculated alone on leaves or co-inoculated with the wild-type strain (2). This suggests that the *syfA* mutant strain did not share in any benefits of syringafactin production by the wild-type strain. Therefore, syringafactin seems to largely affect only the local environment of the cell that produces it rather than serve as a “public good”. Given that bacterial cells on the leaf surface may need only a small, localized quantity of water to avoid desiccation stress (26), the production of a non-diffusible hygroscopic material such as syringafactin might be an economical way for cells to modify their local water environment. In fact, it has been shown that hygroscopic salts on the leaf surface can readily absorb water even at a relatively low ambient RH (26), suggesting that the same phenomenon can occur for syringafactin.

These studies of the water available to bacteria colonizing the apoplast of bean provided great insight into the important role of syringafactin in this habitat and the nature of the plant apoplast itself. The leaf apoplast has been previously described as “a large, air-filled intercellular space” (4). The amount of free water available in the apoplast is still largely unknown (6), but it has been suggested that the apoplast is in fact a dry environment especially when stomata are open (27). Indeed, studies using a *proU:inaZ* reporter gene had indicated that liquid water is apparently largely absent from the apoplast (28). These earlier studies however had not provided any insight as to what the RH might be in the apoplast. It could however be speculated that the water within plant cells would be in close equilibrium with that of water vapor in the apoplast, given the relatively little ventilation that would be expected from diffusion through the stomata. It would be in such a setting that one might expect syringafactin to effectively contribute to the fitness of *P. syringae* since its ability to bind water is much higher in atmospheres that are nearly saturated with water vapor. It was important to note therefore that in the apoplast, the wild-type strain experienced less water stress than the *syfA* mutant at 24 h, and especially at 48 h, after infiltration (Fig. 7B and 7C). Given that *P. syringae* strain B728a is a pathogen of the bean variety used in the study, it was somewhat surprising to find that at least some degree of water limitation was experienced by some cells in both the wild-type and *syfA* mutant strains only 24 h after inoculation (Fig. 7B). A recent study has shown that certain effectors such as HopM1 in pathogens, such as *P. syringae* pv. *tomato* strain DC3000, mediate the release of water from the plant into the apoplast (8). Indeed, at least transient water soaking is a typical symptom of the infection of many plants by pathogenic bacteria. It is thought that the release of water makes apoplastic nutrients more available to bacteria within this habitat and that, since nutrient limitation probably limits bacterial population sizes in the apoplast, water availability is a determinant of the success of a pathogen. In such a setting, it was somewhat surprising that a portion of cells from the wild-type *P. syringae* B728a strain saw lower water availability 24 h after inoculation than in broth media itself (Fig. 7A and 7B). It is possible that effector-mediated water release in bean is transient. However, it is noteworthy that the apoplast becomes even drier between 24 h and 48 h post-inoculation (Fig. 7C). While examining incompatible interactions of plant pathogenic bacteria and host plants, Freeman and Beattie (29) found that plants can actively withhold water from bacterial pathogens that enter the leaf apoplast by 24 h post-inoculation. This suggests that plant responses to the presence of a compatible pathogen such as strain B728a may be delayed and that such water withholding would occur only later during the interaction. The finding that cells of the wild-type strain, and particularly the *syfA* mutant strain, exhibited substantial water stress 48 h after inoculation is consistent with such a model. Earlier work has also shown that hosts such as bean that are compatible with *P. syringae* produce defensive phytoalexins 2 or more days after the infection process is initiated (30). Furthermore, the growth of strain B728a in bean slows with time and typically ceases by 2 days after inoculation (31). This is consistent with a model of decreasing water availability during the infection process. In such a setting, alleviation of water stress by syringafactin production would benefit *P. syringae*.

The demonstration that syringafactin helps provide water to *P. syringae* in natural habitats provides support for an important new role for microbial biosurfactants. It seems likely that at least some of the many microorganisms that produce biosurfactants (19, 32) could similarly benefit. This might be particularly true of those that live in non-water saturated environments, such as soil, that experience periodic water stress. By better understanding the roles of various biosurfactants produced by bacteria, we can gain more insight into the behavior of biosurfactant-producing microbes and the contribution of such compounds to plant-microbe interactions. We should also gain more insight into the interactions between bacteria and the leaf surface. Since biosurfactant producers occur on edible plants, such as lettuce (2), they may influence the behavior of human pathogens, such as *Salmonella*, which can coexist with and benefit from interactions with other epiphytic bacteria (33). Hence, a better understanding of biosurfactant production, and the use of biosurfactant-producing bacteria as biocontrol agents, may help to mitigate both human and plant pathogens thereby improving both human health and crop yield (32).

## MATERIALS AND METHODS

### Bacterial strains and growth conditions

*Pseudomonas syringae* B728a strains were either grown on King’s B medium (KB) plates containing 1.5% technical agar or on half-strength 1/2-21C medium plates (1, 34, 35). Antibiotics were used at the following concentrations (µg/ml): spectinomycin (100), kanamycin (50) and tetracycline (15).

### Syringafactin extraction

Syringafactin was extracted using a protocol described by Burch *et al*. (2) which was modified from a protocol by Berti *et al.* (21). *P. syringae* B728a strains were grown on agar plates for 3 days. Cells were then suspended in water and centrifuged at room temperature at 5000 x g for 10 min. The centrifuge model was a Beckman J2-21M High Speed Centrifuge with a JA-20 fixed angle rotor holding 8 x 50 ml tubes. The supernatant was mixed in a separatory flask with ethyl acetate (1.5:1). The organic fraction was retained while the aqueous fraction was discarded. The organic fraction was reduced to dryness in a rotary evaporator (BÜCHI). The remaining powder was re-suspended in methanol and dried to completion in a Speedvac (Savant).

### Measuring water absorption by syringafactin

Dried, purified syringafactin was added to pre-weighed 1.5 ml microcentrifuge tubes and weighed on an analytical balance (Fisher Scientific) and then placed, with the cap open, in a sealed chamber (magenta box) containing a given saturated salt solution to maintain a constant RH in the air in the chamber (36). Salts that were used were magnesium nitrate, sodium chloride, ammonium sulfate, potassium chloride, potassium nitrate, and potassium sulfate which maintained RHs of 52%, 75%, 80%, 84%, 93%, and 97% respectively. Open tubes were left in the chamber for 3 days before being taken out, rapidly sealed, and reweighed.

### Transformation of *P. syringae* B728a strains

Wild-type *P. syringae* B728a and a *syfA* mutant (31) were both transformed with plasmid pPProGreen carrying a fusion of *proU* with a promoterless *gfp* reporter gene (1) using previous methods (37).

### Preparation of cultured cells for in vitro GFP measurements

Wild-type and *syfA* mutant strains harboring the *proU:gfp* reporter gene were harvested with a loop from half-strength 1/2-21C agar plates that had grown for 1 day at 20°C, and were suspended in 1 ml of half-strength 1/2-21C broth to an adjusted concentration of 10^8^ cells/ml. In addition, in some studies *syfA* mutant cells were suspended to a concentration of 10^8^ cells/ml in 500 µl of half-strength 1/2-21C broth containing 2.5 mg of purified syringafactin. 10 µl drops of each treatment were then pipetted onto 0.4 µm pore-size Isopore^®^ filters that were placed on the surface of half-strength 1/2-21C agar plates. All filters were left on the plates for 8 h at room temperature. After 8 h, filters were transferred to glass slides in chambers containing saturated solutions of magnesium nitrate or water that maintained an atmosphere within the container of 52% RH or 100% RH respectively. After 4 h, filters were placed into 1.5 ml microcentrifuge tubes containing 1 ml of half-strength 1/2-21C liquid to ensure that cells had the opportunity to resuscitate from desiccation stress and translate the *gfp* reporter gene. After 2 h, the tubes containing the filters were vortexed for 20 seconds to remove cells and a 5 µl aliquot of each solution was then pipetted onto a glass slide and left to dry for 20 min. Coverslips were applied to the slides using Aqua PolyMount (Polysciences, cat#18606). Slides were then examined with an epifluorescence microscope as below. It should be noted that the experiments at 52% RH were performed on a different day than the experiments at 100% RH. Thus, the GFP intensity between these experiments cannot be directly compared.

Cell viability assays over time on filters were performed in a similar manner except that filters were exposed to 52% RH as above for various times before filters were removed. Appropriate serial dilutions of cells washed from filters as above were plated onto a KB plate. All plates were incubated at 20°C and colonies enumerated after 2 days.

### Preparation of epiphytic cells for GFP measurements

Wild-type and *syfA* mutant strains were harvested with a loop from half-strength 1/2-21C agar plates grown for 24 h at 20°C and were suspended in 50 ml of water and adjusted to a concentration of 10^6^ cells/ml. Cells were then sprayed onto the leaves of 2-week-old plants of bean (*Phaseolus vulgaris* cv. Bush Blue Lake 274). 4 to 6 seedlings were grown in each pot. All plants were incubated in sealed plastic tents or in large sealed plastic tubs under varying RH conditions maintained with saturated salt solutions as above. Plants were sprayed to wetness and then immediately placed into the sealed plastic tents at 100% RH or into a sealed plastic tub maintained at 97% RH that was then placed in the plastic tent maintaining 100% RH. Plants were kept in the chambers for 2 days to ensure equilibrium. When leaves were to be dried in between chamber transfers, plants were placed at room RH for 20 min to 1 h until water droplets on the leaves had evaporated. Primary leaves were excised from plants (3 leaves per replicate) and were immersed in 150 ml of 5 mM KPO_4_ buffer (pH 7.0) in a beaker. Each beaker was placed in a sonicator (Branson 5510MT) for 10 min to remove cells from the leaf surface. Buffer containing the released cells was filtered through a 0.4 µm pore size Isopore^®^ filter to capture and immobilize the cells. Filters were attached to glass slides and coverslips with Aqua PolyMount. Slides were then examined with an epifluorescence microscope.

### Preparation of apoplastic samples

Wild-type and *syfA* mutant strains were harvested with a loop from half-strength 1/2-21C agar plates that had grown for 1 day at 20°C. Each strain was suspended in 1 L of water to a concentration of 10^6^ cells/ml. Cells were vacuum infiltrated into the leaves of 2-week-old plants (*Phaseolus vulgaris* cv. Bush Blue Lake 274, with 4 to 6 seedlings per pot) as in other studies (9, 30). All plants were stored on the bench at room RH (ca. 50% RH). At time 0, 5 µl of each inoculum was pipetted onto a glass slide and left to dry for 20 min. Coverslips were applied to the slides using Aqua PolyMount. Slides were then used for examination of cells under the epifluorescence microscope. At 24 h and 48 h, primary leaves were excised from plants (3 leaves per replicate) and cut into strips before being immersed in 45 ml of 10 mM KPO_4_ contained in Falcon™ 50 ml conical tubes. Each tube was sonicated for 10 min. After sonication, each tube was also vortexed for 20 seconds. The cell suspensions released from cells was then decanted from each tube into 50 ml centrifuge tubes and centrifuged at room temperature at 7000 rpm (4720 x g) for 10 min. The centrifuge model was Fisherbrand™, accuSpin™ Micro 17 with a 24 x 1.5/2.0 ml rotor. The supernatant was discarded, and the remaining pellets were resuspended in 10 µl of 10 mM KPO_4_. 5 µl of each solution was then pipetted onto a glass slide and left to dry for 20 min. Coverslips were applied to the slides using Aqua PolyMount. Slides were then examined with an epifluorescence microscope.

### Quantification of GFP fluorescence

An M2 AxioImager was used for all microscopic analyses. A GFP filter set was used to view cells in all experiments at 100x magnification and all images were taken in black and white with a 12-bit Retiga camera. Bitplane Imaris image processing and manipulation software was used to quantify the average GFP fluorescence exhibited by each individual cell in digital images. The location of each individual cell was automatically assigned to each cell in a given image, and plant debris and bacterial cellular aggregates were identified visually and de-selected. For each object (individual bacterial cell) identified, the program calculated the mean pixel intensity.

### Statistical Analysis

The software environment R (38) was used to perform the Wilcoxon rank-sum test (39), a nonparametric test of the null hypothesis that it is equally likely that a randomly selected value from one sample will be lesser or greater than a randomly selected value from another sample. The test is appropriate for comparing the distributions of data that are not normally distributed. Mean pixel intensity for each cell from samples was combined and ordered. Ranks were assigned starting with the smallest observation and summed in order to determine the W statistic. R was also used to perform the two-sample t-test (40) to determine if two population means were equal. The test calculates the test statistic, t, and compares it with the distribution of possible values for t to determine a p-value. The two-sample t-test (two-sided) was performed on data that was normally distributed. Statistical significance was determined at a p-value of ≤ 0.05 except for when multiple comparisons were made (Fig. 3). To account for the false discovery rate, the Bonferroni correction (41) was used which resulted in dividing the original p-value of 0.05 by the number of comparisons made. This revealed that each comparison was significant at a p-value of ≤ 0.017.

## ACKNOWLEDGEMENTS

We thank Steven Ruzin and Denise Schichnes at the College of Natural Resource’s Biological Imaging Facility. We also thank Ellen Simms and Fan Dong for their assistance in statistical analysis and Helen Kurkjian for her assistance in statistical analysis and R. This work was supported by the National Science Foundation Louis Stokes Alliances for Minority Participation Bridge to the Doctorate Fellowship, the Chancellor’s Fellowship for Graduate Study, and the William Carroll Smith Fellowship.

## REFERENCES

1. Axtell CA, Beattie GA. 2002. Construction and characterization of a *proU-gfp* transcriptional fusion that measures water availability in a microbial habitat. Appl Environ Microbiol 68:4604–4612.

2. Burch AY, Zeisler V, Yokota K, Schreiber L, Lindow SE. 2014. The hygroscopic biosurfactant syringafactin produced by *Pseudomonas syringae* enhances fitness on leaf surfaces during fluctuating humidity. Environ Microbiol 16:2086–2098.

3. Lindow SE, Brandl MT. 2003. Microbiology of the phyllosphere. Appl Environ Microbiol 69:1875–1883.

4. Melotto M, Underwood WR, He SY. 2008. Role of stomata in plant innate immunity and foliar bacterial diseases. Annu Rev Phytopathol 46:101–122.

5. Remus-Emsermann MNP, Schlechter RO. 2018. Phyllosphere microbiology: at the interface between microbial individuals and the plant host. New Phytol 218:1327–1333.

6. Beattie GA. 2011. Water relations in the interaction of foliar bacterial pathogens with plants. Annu Rev Phytopathol 49:533–555.

7. Monier J-M, Lindow SE. 2003. Differential survival of solitary and aggregated bacterial cells promotes aggregate formation on leaf surfaces. Proc Natl Acad Sci U S A 100:15977–15982.

8. Xin XF, Nomura K, Aung K, Velasquez AC, Yao J, Boutrot F, Chang JH, Zipfel C, He SY. 2016. Bacteria establish an aqueous living space in plants crucial for virulence. Nature 539:524–529.

9. Yu X, Lund SP, Scott RA, Greenwald JW, Records AH, Nettleton D, Lindow SE, Gross DC, Beattie GA. 2013. Transcriptional responses of *Pseudomonas syringae* to growth in epiphytic versus apoplastic leaf sites. Proc Natl Acad Sci U S A 110: e425–e434.

10. Ferro DN, Southwick EE. 2018. Microclimates of small arthropods: Estimating humidity within the leaf boundary layer. Environ Entomol 13:926–929.

11. Drake BG, Raschke K, Salisbury FB. 1970. Temperature and transpiration resistances of *Xanthium* leaves as affected by air temperature, humidity, and wind speed. Plant Physiol 46:324–330.

12. Martin TA, Hinckley TM, Meinzer FC, Sprugel DG.1999. Boundary layer conductance, leaf temperature and transpiration of *Abies amabilis* branches. Tree Physiol 19:435–443.

13. Parlange JY, Waggoner PE, Heichel GH. 1971. Boundary layer resistance and temperature distribution on still and flapping Leaves: I. Theory and laboratory experiments. Plant Physiol 48:437–442.

14. Parlange JY, Waggoner PE. 1972. Boundary layer resistance and temperature distribution on still and flapping leaves: II. Field experiments. Plant Physiol 50:60–63.

15. Schuepp PH. 1993. Tansley review No. 59. Leaf boundary layers. New Phytol 125:477–507.

16. Waggoner PE. 1963. Microclimate and plant disease. Annu Rev Phytopathol 3:103–126.

17. Longrée K. 1939. The effect of temperature and relative humidity on the powdery mildew of roses. Ithaca, N.Y.: Cornell University.

18. Burch AY, Browne PJ, Dunlap CA, Price NP, Lindow SE. 2011. Comparison of biosurfactant detection methods reveals hydrophobic surfactants and contact-regulated production. Environ Microbiol 13:2681–2691.

19. Ron EZ, Rosenberg E. 2001. Natural roles of biosurfactants. Environ Microbiol 3:229–236.

20. Bunster L, Fokkema NJ, Schippers B. 1989. Effect of surface-active *Pseudomonas* spp. on leaf wettability. Appl Environ Microbiol 55:1340–1345.

21. Berti AD, Greve NJ, Christensen QH, Thomas MG. 2007. Identification of a biosynthetic gene cluster and the six associated lipopeptides involved in swarming motility of *Pseudomonas syringae* pv. tomato DC3000. J. Bacteriol 189:6312–6323.

22. Schreiber L. 1996. Wetting of the upper needle surface of *Abies grandis*: influence of pH, wax chemistry and epiphyllic microflora on contact angles. Plant Cell Environ 19:455–463.

23. Yarwood CE, Hazen WE. 1944. The relative humidity at leaf surfaces. Am J Bot 31:129–135.

24. Gnanamanickam SS, Immanuel JE. 2006. Epiphytic bacteria, their ecology and functions, p 131–153. In Gnanamanickam SS (eds), Plant-associated bacteria. Springer, Dordrecht.

25. Leveau JH, Lindow SE. 2001. Appetite of an epiphyte: quantitative monitoring of bacterial sugar consumption in the phyllosphere. Proc Natl Acad Sci U S A 98:3446–3453.

26. Burkhardt J, Hunsche M. 2013. “Breath figures” on leaf surfaces—formation and effects of microscopic leaf wetness. Front Plant Sci 4:422.

27. Zhang D, Tian C, Yin K, Wang W, Qiu J-L. 2019. Postinvasive bacterial resistance conferred by open stomata in rice. Mol Plant Microbe Interact 32:255–266.

28. Wright CA, Beattie GA. 2004. *Pseudomonas syringae* pv. tomato cells encounter inhibitory levels of water stress during the hypersensitive response of Arabidopsis thaliana. Proc Natl Acad Sci U S A 101:3269–3274.

29. Freeman BC, Beattie GA. 2009. Bacterial growth restriction during host resistance to *Pseudomonas syringae* is associated with leaf water loss and localized cessation of vascular activity in *Arabidopsis thaliana*. Mol Plant Microbe Interact 22:857–867.

30. Lyon FM, Wood RKS. 1975. Production of phaseollin, coumestrol and related compounds in bean leaves inoculated with *Pseudomonas* spp. Physiol Plant Pathol 6:117–124.

31. Wilson M, Hirano SS, Lindow SE. 1999. Location and survival of leaf-associated bacteria in relation to pathogenicity and potential for growth within the leaf. Appl Environ Microbiol 65:1435–1443.

32. Burch AY, Shimada BK, Mullin SWA, Dunlap CA, Bowman MJ, Lindow SE. 2012. *Pseudomonas syringae* coordinates production of a motility-enabling surfactant with flagellar assembly. J Bacteriol 194:1287–1298.

33. Poza-Carrion C, Suslow T, Lindow S. 2013. Resident bacteria on leaves enhance survival of immigrant cells of *Salmonella enterica*. Phytopathol 103:341–351.

34. Halverson LJ, Firestone MK. 2000. Differential effects of permeating and nonpermeating solutes on the fatty acid composition of *Pseudomonas putida*. Appl Environ Microbiol 66:2414–2421.

35. King EO, Ward MK, Raney DE. 1954. Two simple media for the demonstration of pyocyanin and fluorescin. J Lab Clin Med 44:301–307.

36. Jarrett DG. 1995. Constant temperature and humidity chamber for standard resistors, p 501–506. In National Conference of Standards Laboratories. Workshop and Symposium (ed), The impact of metrology on global trade: 1995 workshop and symposium. Proceedings of the National Conference of Standards Laboratories, Boulder, CO.

37. Burch AY, Finkel OM, Cho JK, Belkin S, Lindow SE. 2013. Diverse microhabitats experienced by *Halomonas variabilis* on salt-secreting leaves. Appl Environ Microbiol 79:845–852.

38. R Core Team. 2013. R: A language and environment for statistical computing. R Foundation for Statistical Computing, Vienna, Austria. URL https://www.R-project.org/.

39. Wilcoxon F. 1945. Individual comparisons by ranking methods. Biometrics 1:80–83.

40. Student. 1908. The probable error of a mean. Biometrika 6:1–25.

41. Bonferroni CE. 1936. Teoria statistica delle classi e calcolo delle probabilità. Pubblicazioni del R Istituto Superiore di Scienze Economiche e Commerciali di Firenze 8:3–62.

