## Supplemental Figures for "*Pseudomonas syringae* Increases Water Availability in Leaf Microenvironments via Production of Hygroscopic Syringafactin"

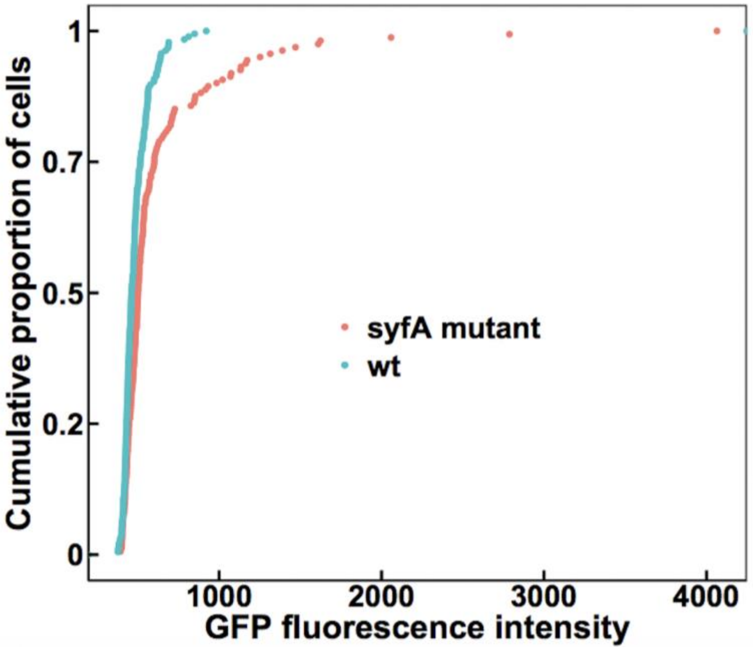


**FIG S1** Wild-type cells and *syfA* mutant cells experience water stress when exposed to 50% RH immediately after inoculation to the leaf surface. GFP fluorescence exhibited by wild-type *P. syringae* strain B728a (blue) or a *syfA* mutant strain (red) harboring a *proU:gfp* reporter when recovered from the leaves of plants incubated at 50% RH for 20 min followed by 100% RH for 2 days followed by drying at 50% RH for 20 min before re-exposure to 100% RH for 2 h. The median fluorescence of wild-type cells was 461 (n = 186) while that of *syfA* mutant cells was 502 (n = 161). Single-cell fluorescence was quantified by microscopy. Significance between sample distributions was tested using the Wilcoxon rank-sum test (wt vs *syfA* mutant: W = 19,168, p-value = $6.76 \times 10^{-6}$).

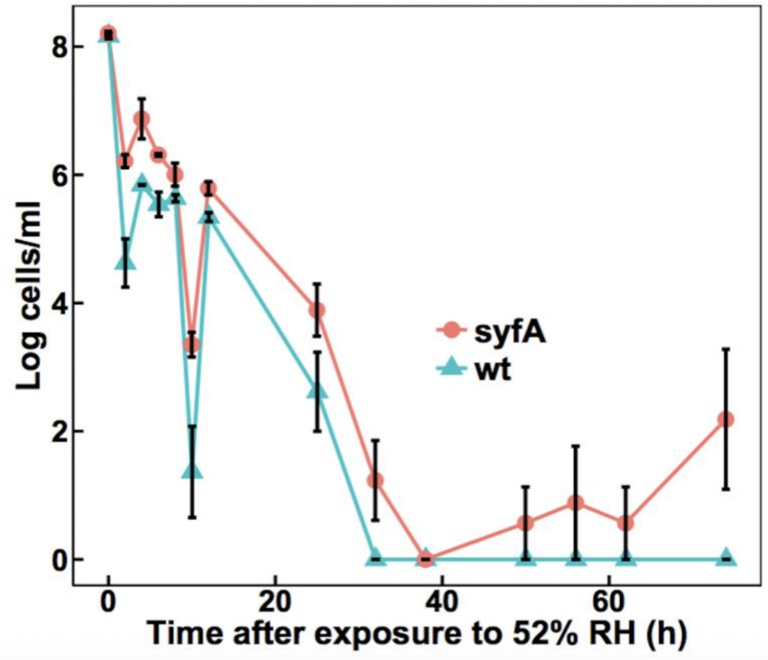


**FIG S2** Proportion of living cells when exposed to 52% RH. Cells of the wild-type, and *syfA* mutant *P. syringae* B728a cells were exposed to 52% RH on filters. The vertical bars represent the standard error of the mean cell concentration.
